# Pre-saccadic enhancement of target stimulus motion influences post-saccadic smooth eye movements

**DOI:** 10.1101/2022.10.10.511640

**Authors:** Shanna H. Coop, Garrett W. Crutcher, Yonatan T. Abrham, Amy Bucklaew, Jude F. Mitchell

## Abstract

Primates move their eyes 2-3 times per second to bring objects of interest to central, high-resolution vision. For moving objects, they use a combination of rapid saccadic eye movements along with smooth following movements to track targets continuously. Each saccadic eye movement produces perceptual enhancements for the target. And for saccades that are made to moving targets, or stationary apertures that contain motion, there is a smooth post-saccadic following response (PFR) of the target motion (Kwon et al, 2019). This PFR occurs involuntarily even when the motion is task irrelevant and could provide an automatic behavioral read-out of the target’s motion. However, PFR movements are small, so it is unclear how reliable they would be as a trial-by-trial read-out. Here we examined PFR in marmoset monkeys performing a foraging task that requires almost no training and which has been shown to involve pre-saccadic neural enhancements of motion selective responses in visual area MT (Coop et al., 2024). We found that PFR in marmosets is highly consistent with humans and could be used to read-out the target motion. More so, we found that the motion in non-target apertures also influenced PFR but to a lesser extent than the target. The gain of PFR was distributed equally between target and non-target apertures before the saccade, and then was rapidly enhanced only for the target motion in driving the post-saccadic following. Thus, PFR provides a behavioral measure of target enhancement relative to distracters in addition to providing a read-out of the target’s motion.

**Significance Statement:** Using a saccade foraging paradigm in marmoset monkeys we measured visual motion integration and pre-saccadic enhancement based on the smooth following eye movements made after a saccade to a motion aperture. We find that marmosets exhibit post-saccadic following behaviors akin to humans, underscoring the evolutionary continuity in visual processing across primates. This following response provided an estimate of target motion that was half as accurate as explicitly trained reports but required minimal training to achieve. It also provided an estimate of the pre-saccadic enhancement for the target relative to distracters in the visual field.

## Introduction

Our visual system processes a wealth of visual information, utilizing saccadic eye movements to ensure high acuity across the visual field. This involves directing our fovea, the area of highest acuity, towards objects of interest, and when those objects are moving to track them. Even before the eyes move, perceptual enhancements occur for the target of eye movements, known as pre-saccadic attention or enhancement (Henderson et al. 1989, Kowler, Anderson, Dosher, & Blaser, 1995; Deubel & Schneider, 1996; Rolfs et al., 2011; White et al., 2013). Pre-saccadic enhancements in human observers have been shown to improve orientation and contrast sensitivity for the targets (Rolfs & Carrasco, 2012; Ohl, Kuper, & Rolfs, 2017), target motion features (White et al., 2013), and the sensitivity for high spatial frequency features (Li, Barbot, & Carrasco, 2016). These enhancements are rapid, occurring roughly 50-100 ms before the saccade (Deubel, 2008), and obligatory, happening even when it impairs task performance (Deubel, 2008; Montagnini & Castet, 2007).

Pre-saccadic enhancement also boosts smooth pursuit eye movements for the motion of the saccade target. Smooth pursuit is used by observers to keep moving objects stabilized near the fovea for continued inspection and better acuity. The initiation of pursuit typically involves a saccadic eye movement to select the target from background and can be followed with other catch-up saccades to correct for slip during tracking (Lisberger, 2010). After a saccade selects a moving target, the post-saccadic smooth eye movements quickly align with its motion excluding competing distracter motion (Gardener and Lisberger, 2001). Recently a similar enhancement for motion of saccade targets was found in humans even when voluntary pursuit was not engaged, termed the post-saccadic following response or PFR (Kwon, Rolfs & Mitchell, 2019). In that study eye movements were cued towards a stationary aperture that contained motion but without any task to track it. Nonetheless, a reliable post-saccadic following response (PFR) was observed which had a low gain along the direction of motion within the stationary aperture. This response persisted even if the stimulus disappeared in saccade flight, demonstrating that it was driven by pre-saccadic enhancement of the target’s motion rather than by passive following of the motion at the fovea in the post-saccadic period. Because this PFR occurs automatically it could offer a simple analog read-out of the target motion, which could be valuable in animal studies as a behavioral report that does does not require extensive training. However, the trial-by-trial accuracy of the PFR has not been assessed, and given the noise associated with small eye movements, could be of relatively limited value as a motion report.

Here we tested the accuracy of the PFR in the marmoset monkey, a small New World primate. The marmoset monkey has gained interest as a model for studies in visual neuroscience due to its rapid breeding, potential for genetic manipulation, and smooth cortical surface, which facilitates neural imaging and recording (Mitchell et al, 2015). Although it is one of the smallest of non-human primates, it retains high visual acuity with a well-defined foveal specialization (Troilo et al, 1993; Nummela et al., 2017), uses saccades to scan visual scenes (Mitchell et al., 2014), and smooth eye movements to track moving targets (Mitchell et al., 2015; Pattadkal et. al., 2024). However, training marmosets on perceptual tasks has proven more challenging than macaques (Mitchell et al., 2014). One study (Cloherty et al, 2020) showed that marmosets could report the direction of motion for a centrally presented random dot field. After extensive training with over 50-70 daily sessions, marmosets were able to make direction judgments with accuracy comparable to humans. Nonetheless, alternative tasks could be of great value if they can provide similar analog reports with less extensive training, and further if they allow for the flexibility to position the motion stimulus at varied peripheral locations.

We adapted a simple saccade foraging task for marmosets to test if they have PFR similar as humans, and if so, how reliably it might read-out trial-by-trial stimulus motion. This task was used previously to measure pre-saccadic enhancements at the neural level in visual area MT (Coop et al., 2024). We tested whether the saccades in this task evoked PFR and compared the reliability of PFR as a read-out of target motion to that of the former study using explicit perceptual reports (Cloherty et al, 2020). Further, because this task includes distracters with independently selected motion directions, we could also measure their influence on PFR and assess the time-course of pre-saccadic enhancement for the target over the distracters.

## Methods

### Subjects and Surgery

All experimental protocols were approved by the [Author University] Institutional Animal Care and Use Committee and were conducted in compliance with the National Institutes of Health guidelines for animal research. Four adult common marmosets (Callithrix jacchus), three males and one female (ages ranging from 2-8 years) participated in a saccade foraging study with eye tracking. Subjects were housed with a circadian cycle of 12-hour light and 12-hour dark. Subjects were briefly food scheduled (delayed mealtime) with full access to water during early training to learn how to perform fixation tasks, but afterwards had no food restrictions at the time behavioral data was collected.

All subjects were surgically implanted with head-posts to stabilize eye tracking. Two months prior to surgery, subjects were trained to sit in a small primate chair (Lu et al. 2001; Remington et al. 2012; Osmanski et al. 2013; Nummela et al. 2017). After head-implant surgery and a 2–3-week recovery period, they were trained to maintain fixation on a small central point and to perform a visual acuity task (Nummela et al., 2017). Each subject’s vision was corrected to optimize acuity using spherical concave lenses (Optimark Perimeter Lens Set) that were centered 4–5 mm in front of the face. The diopters of the lens used in the present study were - 1.0, -2.5, -1.5, and -2.0 respectively for marmosets 1-4.

### Stimulus presentation and timing

Stimuli were generated using the Psychophysics toolbox (Brainard, 1997; Kleiner et al., 2007; Pelli, 1997) in MATLAB 2015b (MathWorks, Natick, MA) on a windows computer (Intel i7 CPU, Windows 7, 8 GB RAM, GeForce Ti graphics card). They were presented on a gamma-corrected display (BenQ X2411z LED monitor, resolution: 1,920 x 1,080 p, refresh rate: 100Hz, gamma correction: 2.2) that had a dynamic luminance range from 0.5 to 230 cd/m2 at 57 cm in a dark room and viewed under head-restraint. Brightness on the BenQ display was set to 100 and contrast to 50, and additional visual features of the monitor, such as blur reduction and low blue light, were turned off. Gamma corrections were verified with measurement by a photometer. Task events and eye movements are recorded using a Datapixx I/O box (Vpixx technologies) for temporal registration. Matlab code is available online (https://github.com/jcbyts/MarmoV5).

Peripheral visual stimuli consisted of random dot motion fields viewed within a Gaussian aperture (**Fig 1A**). Each aperture contained a field of black dots (each dot with a 0.15 dva diameter with a density of 2.54 dots per visual degree squared) which moved at either 6.75 degrees/sec to match parameters in a previous study with humans (Kwon et al, 2019), or at 12 dva/s degrees/sec to better match previous motion processing studies with marmosets (Cloherty et al., 2020; Coop et al., 2024). Each aperture contained dots moving in one direction (100% coherent), with motion selected independently from 1 of 16 directions that spanned 0 to 360 degrees in steps of 22.5 degrees. Depending on the experimental condition, the dots either had unlimited lifetimes or limited lifetimes of 50ms with asynchronous updating to new random locations with the aperture. In the unlimited lifetime condition, each dot had an infinite lifetime in which the dots were replotted on the opposite edge of the aperture once they crossed its boundary. The dots appeared in black (0.5cd/m^2^) against a grey background (115 cd/m^2^). Stimuli were positioned at 5 degrees eccentric locations with a radius of 2.5 degrees. The contrast of dots decreased from full contrast black at the center of the field to zero at the edge according to a Gaussian envelope with a half-width of 1.67 degrees (σ = 0.83 degrees).

### Eye Tracking

Eye position was acquired either at 220 Hz using an Arrington Eye Tracker with Viewpoint software (Arrington Research) or at 1000 Hz using an Eyelink 1000 Plus eye tracker (SR research). The eye was imaged under infrared light through a dichroic mirror (part #64-472, Edmunds Optics) placed at a 45-degree angle in front of the observer, allowing for a direct view of the pupil. Each marmoset viewed the screen through a spherical concave lens to correct for any identified myopia, as described above. The eye tracker was calibrated using a face detection task described previously (Nummela et al., 2017). Raw horizontal and vertical eye position signals were smoothed offline with a median filter (over 5 samples, 5 ms), and if collected at 220 Hz upsampled to 1000 Hz by linear interpolation, and then convolved with a Gaussian kernel (5ms half width, out to 3 SD, -15 to 15ms) to minimize high-frequency noise. For offline detection of saccadic eye movements, we used an automatic procedure that detected deviations in 2-D eye velocity space (Engbert & Mergenthaler, 2006; Kwon et al., 2019). We computed horizontal and vertical eye velocity by taking the temporal difference of the smoothed eye-position traces. Saccades were then marked by events where the 2D velocity exceeded the median velocity by 10 SD for at least 15ms (Engbert & Mergenthaler, 2006; Kwon et al., 2019) and events merged into a single event if they were separated by less than 5ms. Saccade onset and offset were determined by the first and last time the 2-D velocity crossed the median velocity threshold. Any trials in which an eye blink occurred were flagged by the abrupt change in eye position to a location outside the visible screen and were removed from the analysis.

### Behavioral Task

To study post-saccadic following response (PFR), we had subjects perform a saccade foraging task to peripheral motion apertures (**Fig 1A**). Each trial began with the fixation of a small bullseye (0.3 degree outer radius, 0.5 cd/m^2^ center, 230 cd/m^2^ surround) for a delay uniformly distributed between 0.1 to 0.3 s, presented on a gray background (115 cd/ m^2^). Fixation was maintained within a 1.5 degree radius window around the small spot. After the fixation period, three dot motion apertures appeared in the periphery while the fixation spot was offset. For each trial, the direction of motion was independently sampled for each aperture. The apertures appeared at 5 degrees eccentricity in one of two configurations across blocks of 20 trials. The first configuration had a single target above fixation along the vertical meridian with two other targets in the lower hemi-fields each spaced on a circle 120 degrees apart, while the second configuration was the vertically flipped, thus across blocks sampling 6 peripheral locations. We collected behavioral sessions opportunistically on days when marmosets were not involved in other experiments, with sessions lasting roughly 1 hour.

Overall, this study involved three different experimental conditions to afford comparisons first with a previous human study, and then with two different marmoset experiments. In the first condition, each marmoset participated in a minimum of 4 sessions, ranging from 4 to 8 sessions, where they completed up to 600 trials. The trials used a speed of 6.75 degrees of visual angle per second and unlimited lifetime dots, enabling direct comparisons with a previous human study (Kwon et al., 2019). The second condition featured marmosets experiencing motion at a speed of 12 degrees of visual angle per second comparable to previous neurophysiology study that used limited lifetime dots (Coop et al., 2024). In these sessions, both unlimited and limited lifetime dots were randomly interleaved across trials in order to assess the impact of dot lifetime on PFR. Marmosets performed 6 to 9 sessions, totaling up to 1200 trials per session. In the third condition, three out of four marmoset monkeys were involved and completed 12 to 21 sessions, each consisting of up to 600 trials. These sessions exclusively used limited lifetime dots and varied motion coherence (signal strength) by sampling different dot direction bandwidths (0°, 90°, 180°, 270°, 360°) that were matched to an earlier study with explicit perceptual reports of motion direction (Cloherty et al., 2020).

To obtain liquid rewards marmosets were only required to make a saccade to different peripheral apertures across consecutive trials. From aperture onset marmosets had up to 1.5 seconds to select an aperture by saccade, indicated by the first movement exiting the fixation window landing within a 2.5 degree window around the aperture center. Eye position needed to remain in the aperture for 0.2 s to confirm the saccade endpoint to ensure the choice was intentional. In 50% of the trials the motion stimuli disappeared during saccade flight, identified by the moment the eye left the fixation window, such that in the post-saccadic period no foveal motion was present (**Fig 1A**). Saccades that selected an aperture differing from that selected in the previous trial were rewarded with 5-10 uL of liquid reward (marshmallow juice or Ensure) to encourage foraging. An incorrect choice resulted in a black Gaussian spot that filled the incorrectly selected aperture and no liquid reward.

### Trial criteria

Trials contaminated by micro-saccades, double-step saccades, or post-saccadic correctional saccades were excluded from analyses. We excluded trials with micro-saccades of amplitude greater than 0.5 visual degrees during the fixation hold period. To ensure that the animal had not initiated a saccade prior to stimulus presentation, we also excluded trials where the reaction time of saccade initiation was under 0.1 sec or in which the animal made two smaller saccades to the aperture (double-step) as opposed to one larger saccade. Further we excluded trials if a micro-saccade occurred in the 0.2s after the saccade offset in the selected aperture. Overall, this led to removing 9.9% (+/-3.5% sd.) of the total trials. Thus, subsequent analyses only included accurate single step saccades to the peripheral apertures.

### Eye movement analysis

We examined if smooth eye movements tracked the target motion immediately following the saccade offset in the target aperture. An example 2D eye position trace (in black) is shown from one trial to illustrate the low gain post-saccadic following movements in one aperture (**Fig 1D**). The blue arrow indicates the direction of stimulus motion within the aperture while the epoch from 20-60 ms post-saccade, which we refer to as the “Open Loop” period, are labeled in red and indicate drift along a related direction. The eye position traces shown over time (**Fig 1E**) and velocity traces over time (**Fig 1F**) correspond to the same trial. In order to determine if eye velocity drifted in the same direction as target motion, we projected the 2D eye velocity along the motion axis. The projected eye velocity is shown superimposed on a magnified scale (**in red, Fig 1F**), and though noisy indicates on average a positive post-saccadic drift velocity consistent with following target motion.

To quantify the post-saccadic following on a trial-by-trial basis we examined the net drift along the target motion axis during the epoch from 20ms to 60ms after saccade offset, which we term here the Open Loop epoch. This epoch is chosen to avoid contamination of smooth eye movements in the 20 ms immediately following saccade offset and overshoot, and also to be less than the visual latency for foveally presented motion to drive eye movements. Previous human studies used a longer epoch from 20-100 ms reflecting the slower visuo-motor delays in human participants, whereas in marmosets it has been previously established that ocular following is driven within 60-70 ms of motion onset (Cloherty et al, 2020; Pattadkal et al, 2023). To determine the net drift during the open loop period we fit a linear regression line to the 2D eye positions over that interval, creating a 2D vector that we term the post-saccadic following response (PFR) vector. We then projected a PFR vector (**red arrow, Fig 1G**) onto the vector representing the aperture motion over the same period (**blue arrow, Fig 1G**). Comparing the length of the PFR vector projected onto the motion vector provides an estimate of the PFR gain in the open loop epoch. An alternative measure of the trial-by-trial accuracy in reading out the direction of target motion is the angular error between the PFR and motion vectors (**Fig 1G**).

In preliminary pilot tests we noted that post-saccadic drift contained systematic biases in direction that were specific to the target aperture selected by the saccade independent from stimulus motion within it. These biases were generally small, but varied per subject and could limit the fidelity of PFR direction recovered. To control from any systematic drifts in eye position per landing position we computed the net drift per aperture location, averaged across all task trials and including the different target motions uniformly sampled across direction, and subtracted the next from the trial-by-trial raw PFR vectors to give a rescaled vector (**Fig 1H**). All statistics reported use this “corrected” PFR vector, including angular errors.

The previous study in humans (Kwon et al, 2019) also reported that the offsets in saccade landing positions were deviated along the direction of stimulus motion. To test for this in marmosets we followed the analyses of the previous study and defined a saccade end-point vector, measured as the eye position at the end of the saccade minus the center of the aperture. As with the PFR vector, end-point vectors were projected onto the stimulus motion and were rescaled for any systematic deviations per saccade landing position by subtracting the average end-point vector per aperture location. Therefore, positive deviations in the end-point vector reflect saccades that anticipate target motion.

## Results

We measured post-saccadic smooth eye movements in marmosets to test whether they exhibit following for target motion as previously observed in humans (Kwon, Rolfs, & Mitchell, 2019). Marmosets performed a simple saccade foraging task that required minimal training (**Fig 1A**). In each trial, the marmoset fixated upon a central point for 100-300ms, after which three peripheral apertures appeared each which contained random dot field motion. Each aperture contained 100% coherent moving dots with direction sampled independently from 1 of 16 directions (360 degrees, 22.5 degree increments). Marmosets were rewarded for making a single-step saccade to one of the three apertures, with juice reward if they selected a different aperture across adjacent trials to encourage foraging. The same tasks had been used previously to study pre-saccadic enhancement of neural responses to stimulus motion (Coop et al., 2024). The target apertures were presented in two different spatial configurations across blocks of 20 trials. The first configuration was a triangle with an upwards point and two lower base targets, and the second was flipped vertically.

**Figure 1.**
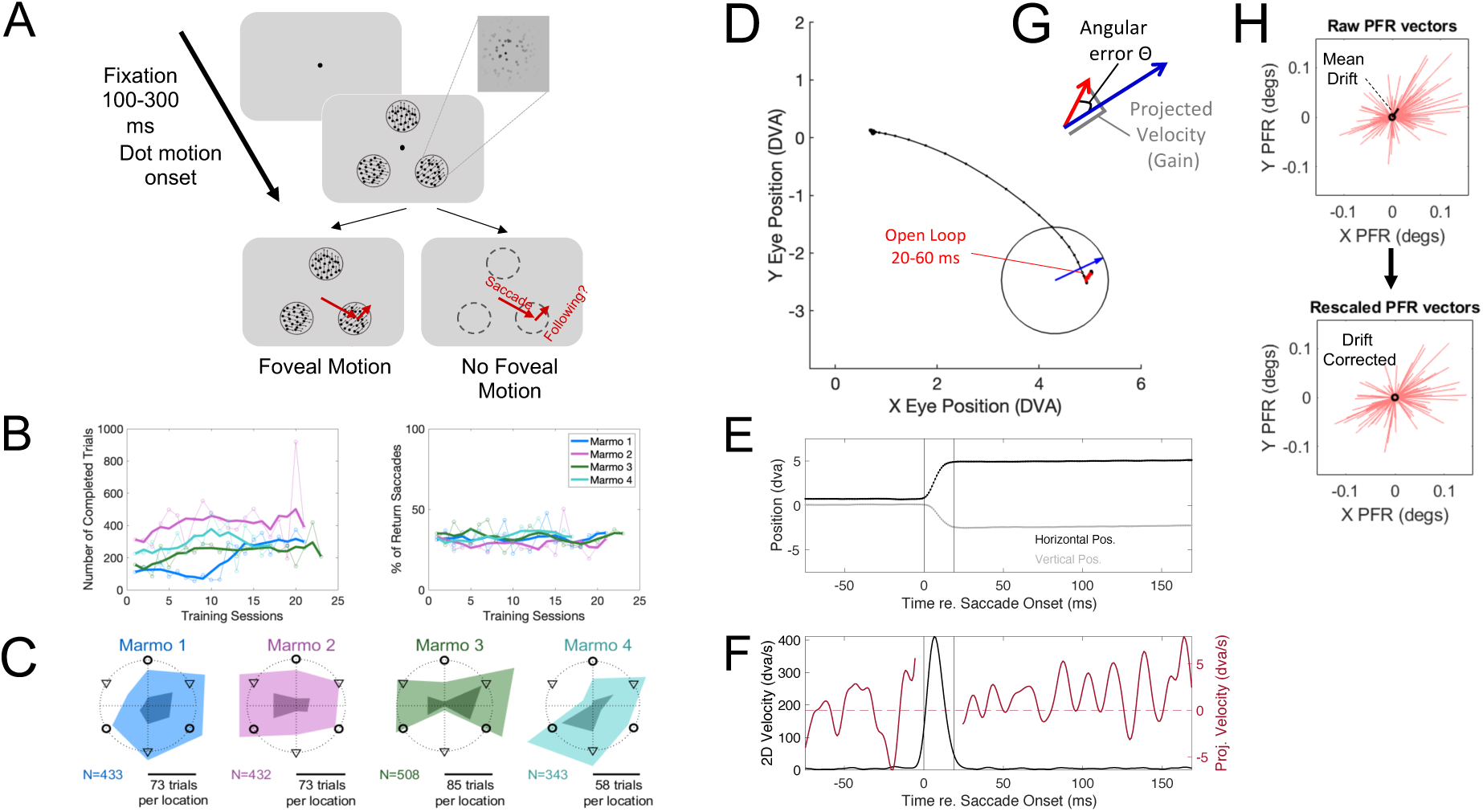
Experimental paradigm and marmoset behavior. (**A)** For each trial, the marmoset held fixation for 100-300ms after which it was presented with three equally eccentric motion apertures. Dot-motion apertures contained 100% coherent motion sampled independently from 16 directions. Each aperture had a Gaussian window overlay. We rewarded saccades to any aperture as long it differed from the previous trial to encourage foraging. In 50% of trials, the dot motion stimuli disappeared during saccade flight such that no stimulus motion was ever presented foveally (i.e., Foveal Motion vs. No Foveal Motion). (**B) (Left)**Total number of completed trials per session increased during the training period after learning a saccade detection task that validated their visual acuity (see methods). Solid lines are the running average across 5 sessions, while open circles represent the raw data. (**Right)** he percentage of return saccades remained stable across the training period session. (**C)** Marmosets exhibited different spatial biases for average trials completed as a function of the size aperture locations (indicated by the lighter filled regions). They also performed a significant fraction of return saccades to a location chosen in the previous trial (indicated by the darker filled regions). Colors match legend in Panel B. (**D)** The 2D traces of eye position from a single trial demonstrate a drift along the target motion during the open loop from 20-60ms following saccade offset. The black dots represent raw eye-position samples from the eye tracker. The eye position began within the fixation window and jumped to the target aperture (indicated by the black open circle). Eye position during the open-loop is indicate in red. Eye position drifts along the same direction as the stimulus motion in the aperture although with reduced amplitude relative to the actual dot displacement of the stimulus (indicated by the blue arrow). (**E)** The raw horizontal and vertical eye position for this example trial (black dots: horizontal, grey dots: vertical) along with the smoothed position traces (lines) are shown. (**F)** The 2-D eye velocity (black line) is shown for this example trial with the interval flagged as a saccade indicated by the black vertical lines. The eye velocity outside the saccade is shown after being projected (dot product) along the target motion direction (red line). Positive velocities indicate that the eye is moving along the same direction as the target motion. (**G)** Two measures of following were computed, the PFR gain which is the projection of the PFR vector (in red) along motion direction and the PFR angular error which is the difference in direction between the PFR and motion vector. The blue arrow depicts the vector for target motion, while the red arrow depicts the PFR vector for this trial. (**H)** We corrected for any mean drift per aperture location that would have biased PFR vectors. The left plot indicates the raw PFR vectors across a single example session in red, with the mean drift potted in black. The right plot indicates the rescaled PFR vectors after subtracting away the mean drift for that location.

Even in the initial training sessions marmosets completed several hundred trials of this task, reflecting that it required minimal training. Marmosets had first learned to hold fixation briefly and make saccades to peripheral apertures to detect Gabor gratings of varying spatial frequency to validate their visual acuity following methods of a previous study (Nummela et al., 2017). We quantified their performance following the introduction of the foraging task. We calculated across training sessions the total number of trials with correctly completed single step saccades and the fraction of saccades that returned to the aperture of the previous trial (return saccades) which received no liquid reward (**Fig 1B**). Marmosets completed on average 202 (std. 87, range 112 to 313) trials in the first exposure to the foraging task and over 20 training sessions increased the average number of trials completed by 25.6% (average trial completed over animals vs. training session: r = 0.73, p = 0.0008). The proportion of return saccades was on average 32.9% (std. 1.6%) in the first session, and though making return saccades resulted in no reward, we nonetheless did not find a significant reduction in their frequency in making them over training (average proportion of return saccades across animals vs training session: r = 0.10, p = 0.69). In brief, marmosets performed the task reasonably well even upon its first introduction and subsequently showed a gradual increase in the number of trials completed over continued training. Of note, these training requirements were modest as compared to a previous study that used perceptual reports of motion (Cloherty et al., 2020). Like the current study, that study first trained marmosets to fixate and perform saccades to peripheral apertures, which required 1-2 months, but then additionally required 50-70 sessions to acquire the task structure and obtain accurate read-out of motion direction.

Once trained, we allowed marmosets to work up to 600 trials using stimuli with unlimited lifetime dots (6.75 dva/sec) for comparison to previous human studies. In those sessions they completed an average of 394 saccade trials (sd. 133). In later sessions their ability to complete larger number of trials continued to improve. In testing conditions including dots of limited lifetime and varying coherence marmosets were allowed to work up to 1200 trials. In those sessions they completed on average 876 (+/-205 sd.) saccade trials. For subsequent analyses of their post-saccadic eye movements, we additionally required that there were no micro-saccades during the brief fixation epoch before the saccade and that saccades to the chosen aperture involved only a single step, with fixation held for 200 ms at the choice (see methods). This removed on average 9.9% (+/-3.5% sd.) of the total completed trials, giving on average of 355 trials in the early sessions with 600 trials and 789 included trials with the 1200 trial sessions.

Marmosets learned to forage between the three apertures with varied location biases. Each marmoset exhibited different average completed trials counts for each of the six possible aperture locations (**Fig 1C, light-filled regions)**. Marmosets 1 and 4 preferred right spatial targets while marmosets 2 and 3 avoided targets at the vertical meridian. And although marmosets were rewarded for selecting a different aperture in consecutive trials, they often made a saccade back to the same location, especially if the target aligned with their spatial bias (**Fig 1C, dark-filled regions)**. We find that on average they made 35.3% (range 32-40) return saccades, a value that did not differ significantly from their first exposure to the task in early training. Further analyses included all trials regardless of whether they involved a relapse to a previous location.

We found that the average eye position tracked the target motion immediately following saccade offset. For an example trial, the 2D eye position drifts after saccade landing along a direction influenced by target motion over the same period (**Fig 1D**). To measure the degree of following in each trial we computed the PFR gain, defined by the shift in eye position during the open-loop period (20-60 ms). The open loop period was defined from 20-60 ms to 1) avoid contamination from saccadic transients within 20ms of saccade offset, and 2) eliminate the influence of post-saccadic motion at the fovea from driving following at a visual delay, which we estimated no shorter than 60ms (see methods). In the example trial shown, the drift in eye position during the open-loop period (shown in red) follows a similar direction as the stimulus motion (indicated by the blue vector). The amplitude of the drift is reduced relative to stimulus motion, reflecting a low 10-20% gain in following. For each trial, we computed the PFR gain by projecting the PFR vector onto the motion vector and normalizing it by the speed. A PFR gain of 1 would indicate matched speed to the target motion whereas a negative value reflects eye velocity in the opposite direction.

We also consider how accurately the PFR vector aligned with the target motion defined by the angular error in each trial (**Fig 1G**). The angle of the PFR vector provides a trial-by-trial estimate of motion direction, with the angular error being closer to zero when PFR is aligned with target motion. One issue with the angular error is that any small systematic drift occurring specific to the saccade landing position, such as systematic drift back to fixation center, will impact the error measure reducing its accuracy. Although small, we did observe at some aperture locations there was a mean post-saccadic drift that was independent of stimulus motion (**Fig 1H**). We corrected for mean drift by computing the average drift across all trials per saccade landing position, with motion direction balanced by being sampled uniformly per location, and then subtracted the mean drift from the PFR vectors. This rescaled PFR vector was used in subsequent analyses for computing gain and the angular error. We also considered how the saccade end-points were offset from the center of the aperture and if they deviated towards target motion. Like the PFR vector, we computed a saccade end-point vector and subtracted off the mean across all trials per aperture location.

### Marmoset monkeys exhibit comparable PFR gain and saccade end-points as humans

We first compared post-saccadic following movements in marmosets to that of humans in a previous study (Kwon et al, 2019). In the first set of experimental conditions, we employed moving random dot fields with unlimited lifetime dots and moving at 6.75 deg/sec that matched stimulus used in the human study. Two remaining key differences were that a line cue instructed humans on where to saccade, whereas marmosets in our study foraged without any explicit cue. A second difference was that in the human study the number of motion directions sampled in each aperture was limited to only two, those moving tangent to the center of the screen (thus clockwise or counter-clockwise). By contrast in the present study, we intended to evaluate if PFR could be used to read-out motion of any direction, not only tangent to the center-out saccade, so we sampled from a full range of directions. To aid in the comparison with the human study, we begin our analyses restricted to cases where stimulus motion was within 45 degrees of tangent to the center-out saccade.

**Figure 2.**
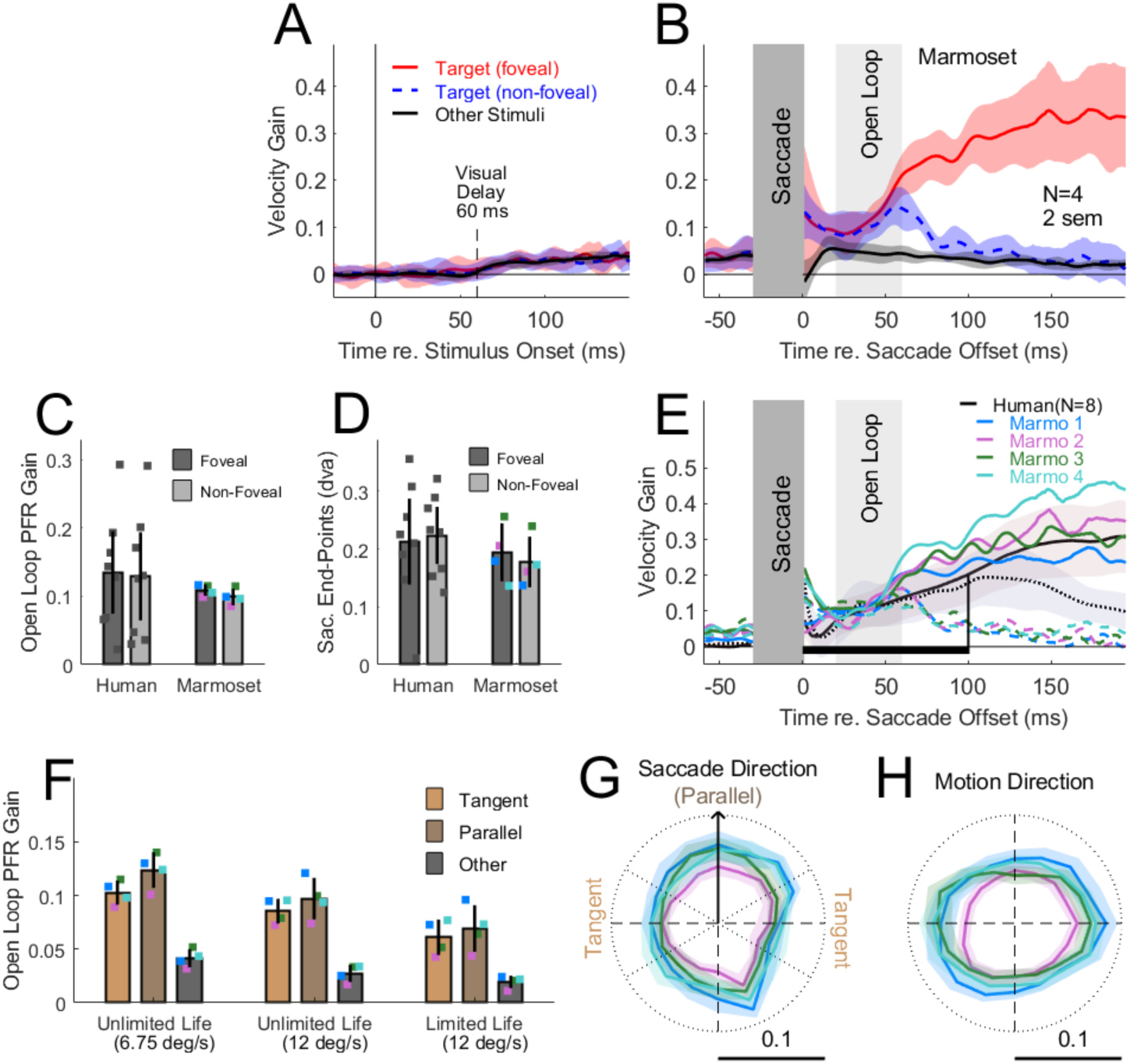
Post-saccadic following response (PFR) and saccade end-points tracked target motion for both humans and marmosets. (**A)** The eye velocity gain projected along the target motion axis is shown averaged across 4 marmosets aligned to the stimulus onset. Positive values reflect following of target motion. Three conditions are superimposed in color: first in red and blue is the following for motion in the target aperture (in red, from trials where the target remains present after the saccade resulting in foveal motion; in blue, where the target disappears in saccade flight, termed non-foveal motion) and the third is following for motion in the non-target apertures (shown in black). Error bars represent 2 SEM across subjects. (**B)** Mean eye velocity traces aligned to the saccade end-points (same conventions as in panel A). We focused subsequent PFR analysis on the open-loop interval from 20 to 60ms after the saccade. The velocity traces during the saccade transient are occluded (shaded in dark gray). (**C)** The PFR gain in the open loop period across humans and marmosets with individual dots representing each participant. Colored dots indicate individual marmosets. Error bars are 2 SEM. (**D)** The saccade end-points along target motion across humans and marmosets (same conventions as in panel C). (**E)** Velocity gain traces over time for individual marmosets (in color) and the mean from humans (N=8, black). Solid traces represent velocity gain when the stimulus is present after the saccade (foveal motion) and dotted traces when absent (no foveal motion). The shaded region marks the open-loop period (20-60ms) and the black horizontal bar the open-loop period in humans (20-100ms). (**F)** Comparison of the PFR gain during the open loop period in marmosets across different speeds and dot lifetime conditions and broken out based on the target motion being tangent (yellow) or parallel (brown) with the center-out saccade, or for the case of non-target aperture motion (black). Error bars are 2 SEM. (**G)** The PFR gain averaged across limited lifetime dots at 12dva/sec are shown as polar plots broken out by the target direction expressed relative to the center out saccade direction (indicated by the upward black arrow). Across marmosets the gain showed a modest bias for larger gains along the axis of the saccade. (**H)** Polar plots of PFR gain broken out by target motion (irrespective of saccade direction) reveals an overall bias for horizontal stimulus motion. Error bars are 2 SEM in G-H.

Even before the saccade the eye velocity exhibited a low gain following response for stimulus motion that began after a visuo-motor delay from stimulus onset (**Fig 2A**). This early drift persisted into the pre-saccadic period and then increased for the target in the post-saccadic period (**Fig 2B).** When time-locked to stimulus onset we find that the average projected velocity shows a modest increase from zero to low-gain following roughly 5% after a visual delay of 60 ms (**Fig 2A**). We examined if this following would occur in relation to other targets in the visual field, the distracters, by projecting the following movements along their axis of motion. This early low-gain response was present for all motion stimuli onset in the visual field, regardless of whether they were the target of the upcoming saccade or a distractor. To visualize the time-course of eye velocity we superimposed the average velocity gain for both the target conditions (red and blue) along with the gain if referenced to non-target stimuli (black). All conditions showed a low 5% gain that was not significantly different when time-locked to stimulus onset (**Fig 2A**), and which persisted into the pre-saccadic epoch 50 ms before the saccade without diverging from each other (**left, Fig 2B**). Thus all stimuli appeared to influence the early drift of the eyes, although with a very low gain.

Immediately after the saccade, PFR gain diverged to favor the target motion over the other aperture motions (**Fig 2B**). The drive for the motion of other apertures remained low and gradually decayed (**Fig 2B, black)**, but for the target motion it abruptly increased in the open-loop period (**Fig 2B, red and blue)**. We further differentiated the following for the target into two types of trials, the first in which the motion stimulus remained present after the saccade (in red: foveal motion) and the second in which the motion stimulus disappeared in saccade flight (in blue: no foveal motion). Immediately after the saccade within the Open-Loop period (20-60 ms), the PFR gain was indistinguishable for target motion regardless if the stimulus remained (red) or disappeared (blue) in saccade flight, reflecting that the presence of foveal motion did not yet make a difference on response due to the visuo-motor delay. However, after the Open-Loop period the velocity gain diverges with the presence of foveal motion driving a stronger PFR gain (red) and otherwise decay without foveal motion (blue). The divergence at 60 ms is consistent with the visuo-motor delay of low gain following seen across the visual field at the initial stimulus onset (**Fig 2A**) as well as previous estimates for ocular following (Cloherty et al, 2020; Pattadkal et al, 2023).

The average PFR gain during the open-loop period indicated a following response for the target of the saccade with 10-15% gain in both marmosets comparable to humans (**Fig 2C**). We projected the PFR vector from the open-loop period onto the target motion for each trial, normalized by target speed, and averaged across trials to estimate PFR gain for each participant. Humans showed an average PFR gain of 13.2% while marmosets showed a 10.4% gain, both which were significantly above zero reflecting drift along the target motion direction (**Table 1**). A two-way ANOVA revealed there was no significant main effects for the difference between foveal versus non-foveal motion, for differences between human versus marmosets, nor was there any interaction (**Table 1**).

Marmosets also exhibited systematic shifts in the saccade end-points along the target motion direction that were similar as humans (**Fig 2D**). We computed the saccade end-point vector, rescaled it for any mean shifts per aperture location, and projected it along target motion (offset in degrees visual angle). Positive deviations reflect the saccade was tilted along the target motion axis. The saccade end-points were on average 0.21 dva in humans and 0.18 dva in marmosets, significantly deviated along target motion (**Table 1**). A two-way ANOVA revealed no significant differences between foveal and non-foveal motion in the saccade end-point deviations, no differences between human versus marmosets, nor any significant interaction (**Table 1**). Thus, both in terms of PFR gain and saccade end-point deviations, marmosets show prediction for target motion in their saccade end-points as well as the post-saccadic smooth following.

**Table 1.** PFR Gain and Saccade Endpoints in Humans and Marmosets.

| **Condition** | **N** | **Open Loop Period (ms)** | **PFR Gain (%)** | **t-value (gain vs zero)** | **p-value** | **Saccade Endpoints (dva)** | **t-value (dva vs zero)** | **p-value** |
| --- | --- | --- | --- | --- | --- | --- | --- | --- |
| Humans | 8 | 20-100 | 13.2 | 4.26 | 0.0037 | 0.21 | 7.37 | 0.00015 |
| Marmosets | 4 | 20-60 | 10.4 | 21.36 | 0.0002 | 0.18 | 8.61 | 0.0033 |
| **Two-Way ANOVA:** |  |  |  |  |  |  |  |  |
|  |  |  | **PFR Gain** |  |  | **Saccade Endpoints** |  |  |
| **Condition** |  |  | **F-value** | **(df<sub>n</sub>,df<sub>d</sub>)** | **p-value** | **F-value** | **(df<sub>n</sub>,df<sub>d</sub>)** | **p-value** |
| Main Effect: Foveal vs non-Foveal Motion |  |  | 0.06 | (1,20) | 0.81 | 0.01 | (1,20) | 0.93 |
| Main Effect: Humans vs Marmosets |  |  | 0.76 | (1,20) | 0.39 | 0.36 | (1,20) | 0.36 |
| Interaction |  |  | <0.01 | (1,20) | 0.94 | 0.16 | (1,20) | 0.69 |

Values are based on data from Kwon et al. 2019 for Humans.

Across individual animals the PFR time-course was highly consistent and comparable to the mean of human observers. The velocity traces per marmoset (**Fig 2E, color traces)** differed modestly in their amplitudes for foveal (solid line) and non-foveal (dashed line) conditions, but overall had consistent qualitative trends. The average from human observers (**black, Fig 2E**) showed a similar qualitative pattern as the marmosets. Both marmosets and human velocity traces were overlapping within the open-loop period for foveal and non-foveal motion conditions, reflecting the presence of a moving dot field in the post-saccadic period was not yet driving following, due to the visuo-motor delay. However, after the open-loop period they diverged, with foveal motion increasing the gain and without it resulting in a decay of following. Overall, humans and marmosets showed a similar time-course with the main difference being the duration of the open-loop period, which in marmosets concluded at 60 ms (shaded area) as opposed to 100 ms in humans (horizontal black line), due to their longer visuo-motor delay.

### PFR generalizes across a full range of motion directions and to limited lifetime dot fields

Having compared against humans, we next considered how PFR generalized to tests including the full range of motion directions. The previous study in humans (Kwon et al, 2019) focused on stimulus motions that were orthogonal to the saccade vector (clockwise or counter-clockwise) in order to reduce the possible impact of saccade transients on the post-saccadic following velocity. While saccades produce a large velocity transient along the direction of the saccade, we have reduced that contamination on the post-saccadic smooth eye velocity by eliminating the first 20 ms after the saccade offset from analyses in our definition of the open-loop period. Thus, it may still be possible to measure PFR reliably even across motions parallel to the saccade and span a full range of 360 degrees. To determine if this was an issue or not, we split the data into halves where aperture motion ran within 45 degrees parallel (yellow) or 45 degrees tangent (brown) to the saccade (**Fig 2F left)**. We pooled the foveal and non-foveal open-loop conditions for this analysis as they did not differ from each other significantly (as seen in Fig 2C). We additionally included the PFR gain in relation to the non-target apertures, which was in all cases modest compared to the target gain (**in dark gray, Fig 2F**). First, we examined the gain for tangent and parallel target motion relative to the saccade direction for the case used to compare against human data, using unlimited lifetime dots moving at 6.75 dva/sec (**left, Fig 2F**). We found that the gain was similar for parallel or tangent motion relative to the saccade direction (12.4% vs 10.2%), and in fact, smaller gain for the tangent than parallel motion case (**Table 2**). The gain for the non-target aperture motion was modest by comparison with only 4.1%, significantly smaller than either tangent or parallel motion for the target (**Table 2**). This suggests that PFR is robust across a range of motion directions and remains boosted for the target in the post-saccadic open-loop period relative to other aperture motions.

**Table 2.** PFR Gain across motion types (Parallel, Tangent, Non-Target)

| Condition | Gain (%) | t-value | df | p-value |
| --- | --- | --- | --- | --- |
| **Unlimited Life (6.75 dva/sec)** |  |  |  |  |
| Parallel Motion | 12.4 |  |  |  |
| Tangent Motion | 10.2 |  |  |  |
| Parallel vs Tangent |  | 6.41 | 3 | 0.0077 |
| Non-Target Motion | 4.1 |  |  |  |
| Tangent vs Non-Target Motion |  | 18.61 | 3 | 0.0003 |
| Parallel vs Non-Target Motion |  | 16.63 | 3 | 0.0004 |
| **Pooled Across Stimulus Types** |  |  |  |  |
| **(Unlimited Life at 6.75 dva/sec, Unlimited Life at 12 dva/sec, Limited Life at 12 dva/sec)** |  |  |  |  |
| Parallel Motion | 8.3 |  |  |  |
| Tangent Motion | 9.6 |  |  |  |
| Parallel vs Tangent |  | 2.94 | 3 | 0.06 |
| Non-Target Motion | 2.9 |  |  |  |
| Tangent vs Non-Target Motion |  | 14.1 | 3 | 0.0007 |
| Parallel vs Non-Target Motion |  | 9.67 | 3 | 0.0023 |

To determine if the speed or dot lifetime made any differences for the PFR gain observed we compared three stimulus conditions: unlimited lifetime dots at 6.75 dva/s and at 12 dva/s, and limited lifetime dots at 12 dva/s (**Fig 2F**). Limited lifetime dots are frequently used in neural studies because they rule out an observer strategy that tracks any single dot, forcing the integration of dot motion to form a coherent percept. A two-way ANOVA on these three stimulus conditions versus motion direction (tangent, parallel, or other) revealed a main effect for the type of stimulus, as well as an effect for motion type, and no significant interaction (**Table 3**). Pooling across stimulus types, there was no significant difference between tangent and parallel target motion with an average gain of 8.3% and 9.6% respectively (**Table 2**), while the gain of the other aperture motion was at 2.9 %, nearly three times smaller (**Table 2**). Between stimulus conditions, the gain was marginally smaller at 12 dva/sec as compared to 6.75 dva/sec (9.1% vs 11.3%) with unlimited lifetime dots (**Table 4**). Comparing unlimited and limited lifetime dots at 12 dva/sec, the limited lifetime dots produced a smaller average PFR gain than unlimited lifetime dots, 6.5% vs 9.1% (**Table 4**). However, the PFR gain was still significant with limited lifetime dots with an average of 6.5% for target motion and lower gain of 1.9% for other aperture motion (**Table 4**). These results show that PFR is robust across stimulus conditions and would be expected with the limited lifetime dots used in previous studies (Cloherty et al, 2020; Coop et al, 2024).

**Table 3.** PFR gain across stimulus (Unlimited Life at 6.75 dva/sec, Unlimited Life at 12 dva/sec, Limited Life at 12 dva/sec) and motion types (Parallel, Tangent, Non-Target)

| Two-Way ANOVA |  |  |  |
| --- | --- | --- | --- |
| Condition | F-value | (df<sub>n</sub>,df<sub>d</sub>) | p-value |
| Main Effect for Stimulus Type | 23 | (2,27) | <0.0001 |
| Main Effect for Motion Type | 74.8 | (2,27) | <0.001 |
| Interaction | 1.39 | (4,27) | 0.26 |

**Table 4.** PFR gain across stimulus types (Unlimited Life at 6.75 dva/sec, Unlimited Life at 12 dva/sec, Limited Life at 12 dva/sec)

| Condition | Gain (%) | t-value | df | p-value |
| --- | --- | --- | --- | --- |
| Unlimited Life (6.75 vs 12 dva/sec) |  |  |  |  |
| 6.75 dva/sec | 11.3 |  |  |  |
| 12 dva/sec | 9.1 |  |  |  |
| 6.75 dva/sec vs 12 dva/sec |  | 3.77 | 3 | 0.0325 |
| 12 dva/sec (Unlimited vs Limited Life) |  |  |  |  |
| Unlimited Life | 9.1 |  |  |  |
| Limited Life | 6.5 |  |  |  |
| Unlimited Life vs Limited Life |  | 9.33 | 3 | 0.0026 |
| Limited Life 12 dva/sec (Target vs Non-Target) |  |  |  |  |
| Target Motion (Parallel + Tangent) | 6.5 | 7.12 | 3 | 0.0057 |
| Non-Target Motion | 1.9 | 6.14 | 3 | 0.0066 |

In order to use the PFR vector to read-out stimulus motion, it should be reliable across a full range of motion directions and not vary substantially based on its alignment (parallel or tangent) with the direction of the saccade performed. We examined the magnitude of PFR when broken out as a function of how target motion aligned with saccade direction using the limited lifetime dot stimuli. Across the four marmosets, we found that PFR gain was largely uniform although there was a bias favoring gain for motion aligned parallel with the saccade direction (**Fig 2G**). We also considered if the magnitude of gain might differ based on the motion of the stimulus itself, independent of saccade direction made to its aperture. Indeed, across the four animals there was a bias for horizontal motions to have higher PFR gain than vertical motion (**Fig 2H**). Despite these biases, the PFR gain was relatively uniform in gain across a full range of directions and relative to saccade direction.

### Trial-by-trial read-out of motion direction by PFR

Whereas the PFR gain measures the drive of motion on eye velocity, we also sought to measure how accurately motion direction can be read-out on a trial-by-trial basis. Therefore, we examined the distribution of angular errors of the open-loop PFR vector, where clustering of errors near zero would indicate alignment with target motion. As illustrated for a single behavioral session, there can be a strong correlation (r = 0.77, circular correlation) between the motion direction of the target and the direction recovered from the PFR vector (**Fig 3A, black)**. We can also compare the trial-by-trial PFR direction against the motion in the other non-target apertures. For the same session we observe a weak correlation (r = 0.05, circular correlation) between the PFR direction and motion direction of the other apertures (**Fig 3B, grey)**. The distributions of angular error reflect tighter clustering around zero for target motion (black) as compared to a broader distribution for non-target, or other aperture, motion (grey) (**Fig 3C**). We fit a maximum-likelihood estimate of a Von Mises distribution to the error distributions (solid lines). Across the four marmosets we averaged the error histograms across sessions for target motion (solid line) and other aperture motion (dashed line) (**Fig 3D**). We observed a consistent pattern in which clustering around zero is narrower for the target aperture motion with a flatter distribution for non-target apertures. Marmosets did differ in the strength of clustering for the target, with marmosets 1 and 4 showing tighter distributions.

**Figure 3.**
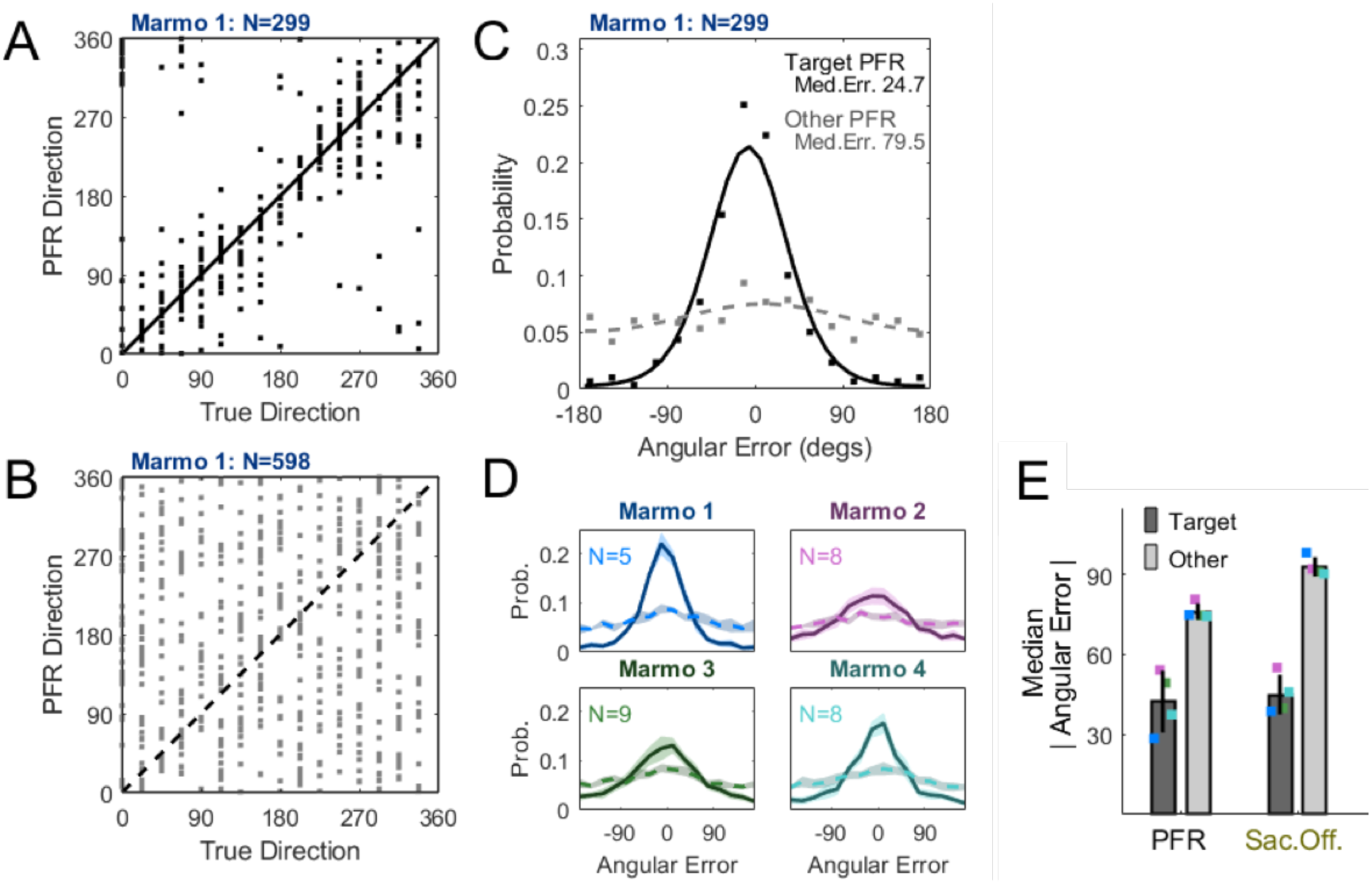
PFR angular error in an example session and across individual subjects. Distributions of angular errors are shown for a single session from Marmoset 1 in A-C. **A)** The direction of motion in the saccade target aperture is plotted against the PFR vector direction (black points indicate individual trials). B) The direction for the other non-saccade targets against the PFR direction is shown (grey points indicate single trials). **C)** The probability histogram of PFR angular errors for the saccade target (black) and the other non-selected apertures (grey-dotted) is shown for the same session. Both distributions were fit with a Von Mises function. **D)** Average angular error distributions from the target (solid lines) and other aperture (dashed lines) for individual marmosets. Error bars are 2 SEM.

How tightly angular error clusters around zero for the target vs non-target apertures provides a measure of the pre-saccadic enhancement of target motion for each animal, and session by session for individual animals. To better quantify the variance across sessions we plotted the median PFR angular error for target motion against that of non-target motion (**Fig 3E**). The median PFR error was 42.6 degrees (+/-11.6 sd.) for target motion averaged across animals as compared to 76.1 degrees (+/-3.0 sd) for other aperture motion. The same analyses can also be applied to saccade end-point vectors (see methods). Like the PFR, target motion drove the saccade end-point vector more than non-target aperture motion. We find that saccade end-points, like PFR, carry more information about target motion than other aperture motion, with a median error of 44.9 degrees (+/ 7.5 sd.) as compared to 92.8 degrees (+/-3.6 sd.). Both PFR and saccade end-points gave comparably accurate measures. In subsequent analyses we focused on how the PFR measure varies with motion signal strength.

### Angular errors from PFR provide roughly half the accuracy of intentional saccadic reports

A previous study examined the distribution of angular errors from saccadic reports made in an analog motion estimation task using limited lifetime dots (Cloherty et al., 2020). They trained two marmosets to view a centrally presented motion dot field at the fovea and make a saccade to a peripheral location on a surrounding ring that extrapolated the central motion. They also varied the signal strength of motion in the dot fields by adjusting the bandwidth of the distribution from which dot directions were drawn, ranging from all dots having a uniform direction (signal strength of 1), to them being drawn over a range of +/-90 degrees around a mean direction (signal strength of 0.5), or completely at random (range of +/-180 degrees giving a signal strength of 0). They quantified the distribution of angular errors in the reported motion direction by fitting an adjusted Gaussian that included a lapse rate across different tested signal strengths (see methods, Cloherty et al, 2020). In three of our four marmosets we were able to perform additional experiments to measure the PFR using stimuli with the same manipulation of signal strengths and fit the angular error distributions using identical methods.

The PFR shows a decreasing error with increasing signal strength that is comparable to saccadic reports but has about half the accuracy (**Fig 4A**). For the explicit perceptual reports, the fit angular standard deviations decreased with increasing signal strength from 0 to 1 reaching a best performance 19.1 (+/-1.7 sem) degrees in angular width (**Fig 4A, black and grey)**. The PFR errors from the three marmosets also decreased with signal strength but only reached a peak accuracy about half as good with 43.4 (+/-2.3 sem) degrees in angular width (**Fig 4A, colored traces)**, which across animals was significantly worse than explicit perceptual reports (**Table 5**). Thus, while PFR provides an automatic read-out for target motion that can be obtained with minimal training, it is, as expected, less accurate.

The second measure of accuracy relates to the proportion of correct choices in a task, which unlike the standard deviation of angular errors in the fit distributions above, also includes errors due to lapses in the task. The previous study (Cloherty et al., 2020) defined correct choices as having an angular error of less than 27 degrees. For explicit perceptual reports they found that percentage correct with signal strength (**solid black and gray lines, Fig. 4B**), whereas percent correct was lower for PFR in the present study (**color lines, Fig 4B**). The peak performance was 78.7% (+/-2.7% sem.) correct for perceptual reports as compared to 37.9% (+/-1.1% sem.) correct for PFR, a significant difference (**Table 5**). Thus, saccadic reports remain more accurate even when lapse rates are included. Of note, the previous study also examined the following responses to foveally presented random dot motion, which they termed “foveal drift” (Cloherty et al, 2020). Like the PFR, the foveal drift was less accurate than explicit perceptual reports made by saccade choices (**dotted lines, Fig 4B**). Foveal drift at the highest signal strengths gave a mean proportion correct of 50.7% (+/-2.6% sem), which was significantly worse than explicit reports (**Table 5**). The foveal drift was still better than the PFR for peripheral stimuli, providing on average 12.8% higher percentage correct than the PFR (**Table 5**). In summary, saccadic reports provide the highest accuracy for reporting trial-by-trial motion, and when comparing different smooth following movements, presentation of stimuli at the fovea rather than periphery (as with PFR at 5 degrees eccentricity) drives following direction more reliably. Nonetheless, the PFR does provide a read-out of target motion that captures a similar dependence on signal strength as explicit reports, and requires minimal training.

**Figure 4.**
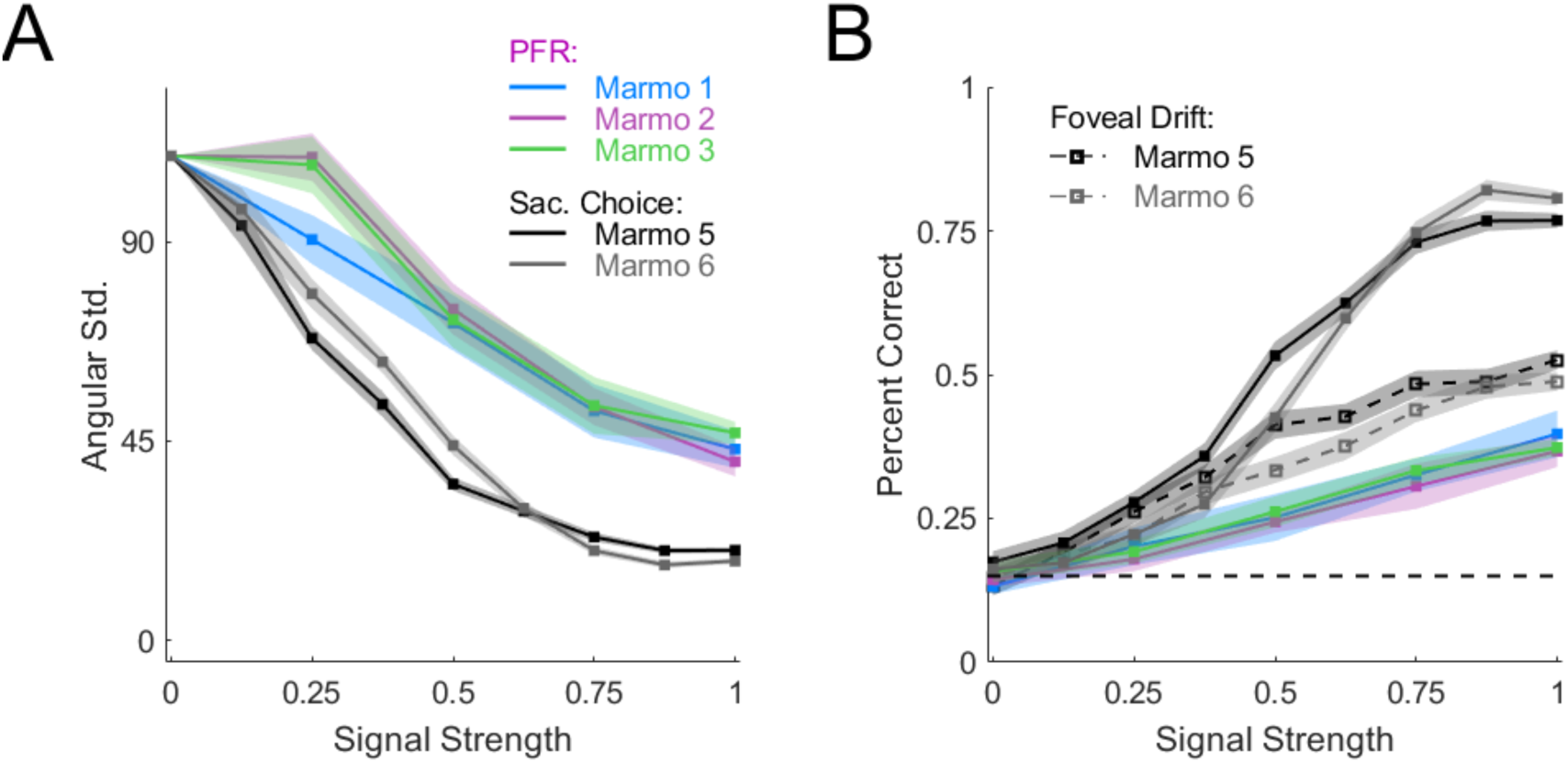
PFR angular errors provide about half the accuracy of a trained analog saccadic choice. **A)** The angular standard deviation narrows across signal strength (motion coherence) for PFR in our three untrained animals (blue, purple, green). By comparison saccadic motion reports are roughly twice as accurate across signal strength (black, grey) in two marmosets (reproduced from Cloherty et al., 2020). Error bars are 2 SEM. **B)** The accuracy of PFR and saccadic choices are replotted from A in terms of task performance with correct choices defined as being within 27 degrees of the true motion direction (same color conventions). In the two marmosets providing saccadic reports (marmosets 5,6) we additionally measured the following response after the onset of a foveal motion stimulus at fixation (Foveal Drift: dotted lines, black, grey and can determine its accuracy as a report (dotted lines, black, grey). Error bars are 2 SEM.

**Table 5.** Accuracy of PFR and Explicit Saccadic Reports.

| Measurement | N | Angular Errors (degrees) | Proportion of Correct Choices |
| --- | --- | --- | --- |
| Explicit Reports | 2 | 19.1 (+/- 1.7 sem) | 78.7% (+/-2.7% sem) |
| Foveal Drift | 2 | (Not reported) * | 50.7% (+/- 2.6% sem) |
| PFR | 3 | 43.4 (+/- 2.3 sem) | 37.9% (+/-1.1% sem) |
| Explicit Reports vs PFR | | $t$ test $p=0.0026$ | $t$ test $p=0.0002$ |
| Foveal Drift vs Explicit Reports | | | Unpaired $t$ test $p=0.0091$ |
| Foveal Drift vs PFR | | | Unpaired $t$ test $p=0.0060$ |

Values are based on data from Cloherty et al. 2020 for explicit reports and foveal drift. \*Angular errors were not reported for foveal drift.

## Discussion

In the current study, we established that like human participants, marmoset saccades to peripheral motion apertures result in smooth post-saccadic following responses (PFR) that tracked stimulus motion at a low gain (Kwon et al. 2019). By including trials in which the stimuli disappeared in saccade flight, we could also isolate the component of post-saccadic following that was due to pre-saccadic motion integration without the influence of foveal motion after the saccade. We found that following did not depend on the presence of foveal motion in the open-loop period (20-60ms), an epoch before post-saccadic visual information can influence the eyes. In marmosets this epoch concluded at 60 ms whereas in humans it extended out to 100 ms, reflecting differences in the species visuo-motor delays. In that open-loop period, we find that following responses are preferentially driven by the motion in the aperture targeted by the saccade with a 10-15% gain, whereas motion from other non-selected apertures has a much weaker influence at or below 5% gain. There was no evidence for preferential weighting of motion in the target aperture before the saccade onset. Rather there was a weak gain (<5%) equally distributed across all apertures in the visual field. The change in weighting of stimulus motion in the open-loop period immediately after the saccade thus reflects the result of a pre-saccadic enhancement of target motion that is then observed immediately in the post-saccadic following. Last we evaluated how accurately PFR provides a trial-by-trial measure of the target motion direction based on the angular error of the following response. Although PFR angular error was about half as accurate as that obtained in a previous study with explicit perceptual reports of motion direction (Cloherty et al., 2020), it does provide a coarse read-out and hold a key advantage in terms of training time.

The current study employed a saccade foraging task which required minimal training. These findings thus demonstrate that the PFR generalizes to less stringent task and stimulus conditions than used in the previous human study (Kwon et al., 2019), where participants were cued where to make a saccade by a flashed line at fixation. Additionally, the former human study tested a set of restricted aperture motions which were tangent to saccade direction, whereas our study extends those results to a full range of motion directions. By restricting our analyses to 20 ms after the saccade offset, we avoid contamination of eye velocity from saccadic transients and minimize the impact of saccadic velocity along the parallel direction. We find that it is indeed possible to measure the PFR across a full range of motion directions, and that gain is equal or slightly stronger for directions parallel to the saccade as compared to tangent to the saccade, and thus not reduced due to saccadic contamination. We also confirmed that the PFR is present with limited lifetime dot stimuli, which are valuable for examining variations in stimulus signal strength and frequently employed in neurophysiological investigations of motion processing.

The PFR likely relies on different neural pathways to read-out motion information than perception. It is well established that smooth eye movements can be disassociated from the perception of motion (Spering and Gegenfurtner, 2007; Spering et. al., 2011; Spering and Carrasco, 2012). Smooth eye movements are more directly influenced by retinal motion whereas perception is influenced by contextual factors. A recent study on the recovery of motion perception in cortically-blind patients with V1 lesions found that improvements in perception did not correlate with any recovery of PFR, even when perception was recovered to normal levels (Kwon et al., 2022). In fact, they found no significant PFR in blind-fields, suggesting it likely requires motion information that passes through area V1 and is lost after V1 lesions. They did, however, find some residual effect for saccade end-points retaining the bias along the direction of target motion, although much weaker than in normal controls. There is reason to suggest saccade end-points may correlate better with perception. Previous work has found that motion can bias the perceived position of peripheral targets, which in turn biases saccades towards them (Schafer and Moore, 2007; Kosovicheva et al., 2014). Further work will be necessary to determine how read-out pathways differ for saccade end-points and PFR.

These experiments show that PFR is sensitive not only to motion in the target apertures, but also from the other non-selected apertures. Thus it can provide behavioral measures of pre-saccadic enhancement of target information relative to non-target apertures. We found that after stimulus onset but before the saccade all apertures in the visual field exerted an equal influence on smooth eye movements, with a weak gain roughly at 5% of aperture motion. However, immediately after the saccade in the open-loop epoch the motion information for the target was boosted, consistent with a post-saccadic enhancement found in voluntary pursuit tasks where a saccade selects between one of two moving targets (Gardener and Lisberger, 2001). Studies of motion processing in area MT of the macaque have shown that variability in neural activity correlates with variations in smooth pursuit behavior (Huang and Lisberger, 2009; Lisberger, 2010). Like pursuit, PFR is a smooth eye movement that provides an analog read-out of motion information and reflects a boost in the gain for target motion during saccade planning. Unlike smooth pursuit tasks and tasks requiring explicit perceptual reports, the PFR occurs without any explicit training, which though it provides a less accurate motion read-out, may prove useful for investigations of the neural mechanisms of motion processing and target selection.

